# Transfer learning improves antibiotic resistance class prediction

**DOI:** 10.1101/2020.04.17.047316

**Authors:** Md-Nafiz Hamid, Iddo Friedberg

**Author notes:** Molecular Pathology Unit and Center for Cancer Research Massachusetts General Hospital Research Institute, Charlestown, USA and Department of Pathology, Harvard Medical School, Boston, USA.

## Abstract

**Motivation:** Antibiotic resistance is a growing public health problem, which affects millions of people worldwide, and if left unchecked is expected to upend many aspects of healthcare as it is practiced today. Identifying the type of antibiotic resistant genes in genome and metagenomic sample is of utmost importance in the prevention, diagnosis, and treatment of infections. Today there are multiple tools available that predict antibiotic resistance class from DNA and protein sequences, yet there is a lack of benchmarks on the performances of these tools.

**Results:** We have developed a dataset that is curated from 15 available databases, and annotated with their antibiotic class labels. We also developed a transfer learning approach with neural networks, TRAC, that outperforms existing antiobiotic resistance prediction tools. While TRAC provides the current state-of-the-art performance, we hope our newly developed dataset will also provide the community with a much needed standardized dataset to develop novel methods that can predict antibiotic resistance class with superior prediction performance.

**Availability:** TRAC is available at github (https://github.com/nafizh/TRAC) and the datasets are available at figshare (https://doi.org/10.6084/m9.figshare.11413302).

## 1 Introduction

Each year at least 2 million people in the US are affected by antibiotic resistance, and at least 23,000 fatally [1]. In the European Union, it is estimated that 30,000 people die each year from antibiotic resistance related causes [2]. Lacking interventions, it is estimated that 10 million people will die from antibiotic resistant infections each year by 2050, and the total GDP loss resulting from antibiotic resistance is estimated to be equivalent to 100.2 trillion USD [3]. Surveillance of antibiotic resistance is an integral aspect of mitigating the clinical and environmental impacts of antibiotic resistance. Therefore, rapid identification of the antibiotic resistance class from clinical and environmental metagenomic samples is critical. Currently, culture based techniques such as Antimicrobial Susceptibility Testing (AST) are used to classify antibiotic resistance which provides important information to devise patient treatment. While these methods are an important part of the clinical setting, they require lab facilities and experts not available at every setting. They also do not work for uncultured bacteria which usually constitute a large part of any complex microbial community [4]. Computational tools coupled with rapid sequencing tools can complement culture based techniques by identifying the antibiotic resistance class from sampled genome sequences.

Naturally produced antibiotics and antibiotic resistance have co-evolved over billions of years [5, 6]. However, the dramatically increased use of antibiotics in clinical and agricultural settings has led to a strong selective pressure and rapid evolution of novel antibiotic resistance genes, and their concurrent acquisition via horizontal gene transfer. Computational tools can help identify these genes along with the antibiotic class they confer resistant to, but there is a dearth of datasets through which we can assess the relative performance of these computational tools.

To address this issue, we compiled COALA (COllection of ALl Antibiotic resistance gene databases) dataset. COALA data were compiled from 15 antibiotic resistance databases available online, and include sequence data labeled for 15 types of antibiotic resistance. We provide three versions of this dataset. (1) COALA100, the full aggregate of all the databases; (2) COALA70 reducing COALA to no more than 70% sequecne ID between all sequence pairs and (3) COALA40, reduction to 40% sequence ID between any sequence pair. The reductions were done using CD-HIT [7].

We then developed a deep neural network based transfer learning approach TRAC (TRansfer learning for Antibiotic resistance gene Classification), that can predict the antibiotic classes to which the genes are resistant to. Current state-of-the-art tools such as CARD-RGI [8] NCBI-AMRFinder [9], and SARGFAM [10] use alignment based methods such as BLAST [11] or HMMER [12] or augmented versions of these methods. Recently, a deep learning based approach, DeepARG [13], was developed that used normalized bit scores as features that were acquired after aligning against known antibiotic resistant genes.

In typical supervised learning approach, an algorithm is trained to solve a task, and is then expected to solve the same task using future unseen data. Supervised learning requires ample labeled data, but it is quite common that in genomic data researchers do not have enough labeled data for specific tasks, where not many genes are known to perform specific functions. In the case of antibiotic resistance spanning 15 classes, there are not enough positive examples populating all classes to perform training and testing. To overcome this problem, we use transfer learning, a machine learning technique where a learning algorithm is first trained to solve a task, and then uses the trained algorithm to solve a related but different task [14]. Transfer learning allows the use of labeled or, more importantly, unlabeled data to train an algorithm, and then train again on a small corpus of labeled data. For example, in any kind of image classification task, it is now commonly accepted to use a pre-trained model trained on a large labeled dataset such as Imagenet [15], and then again train that model on a smaller labeled dataset of the final task (known as fine-tuning) [16, 17]. This is an example where a large corpus of labeled data is being used to augment the performance in a task where labeled data are scarce. Using another example from Natural Language Processing (NLP), it is common to use a pre-trained model that has been trained first as a language model on a large corpus of sentences. These pre-trained models are then used for further downstream supervised classification tasks such as text classification [18]. Language modeling is the task of estimating the probability of a sentence happening in that language [19]. This type of language model is typically trained to predict the next word in a sentence given some words previously. After a language model learns a probability distribution over all possible sentences in a language, the model is trained on a smaller task specific labeled dataset such as sentiment classification e.g. predict from a given review if the review is positive or negative [18].

Transfer learning has been shown to be effective in various tasks of protein sequence classification [20, 21, 22, 23] where there is a small number of labeled data. Currently, there is a large number of unlabeled protein sequences publicly available despite the small amount of labeled data for many specific tasks. For example, Uniprot/TrEMBL [24] has ~158M protein sequences. In the same vein of the NLP example above, this large number of unlabeled protein sequences can be leveraged to learn a language model where a deep neural network can be trained to predict the next amino acid in a protein sequence. Here, we have created an artificial supervised task where the label is the next amino acid event though the dataset is unlabeled in its true meaning. By training this way, the neural network learns the probability distribution of all possible protein sequences. In this paper, we used UniRef50 [25] to train our particular neural network language model, and then used the trained neural network to predict the antibiotic class to which the gene confers resistance, by training on a small labeled dataset where the training data are protein sequences, and true labels are the antibiotics. As we show, TRAC outperforms other methods by a large margin, and performs better than a deep learning method that was trained only on a labeled antibiotic resistance sequence dataset, underscoring the advantage of transfer learning for this task.

**Figure 1:**
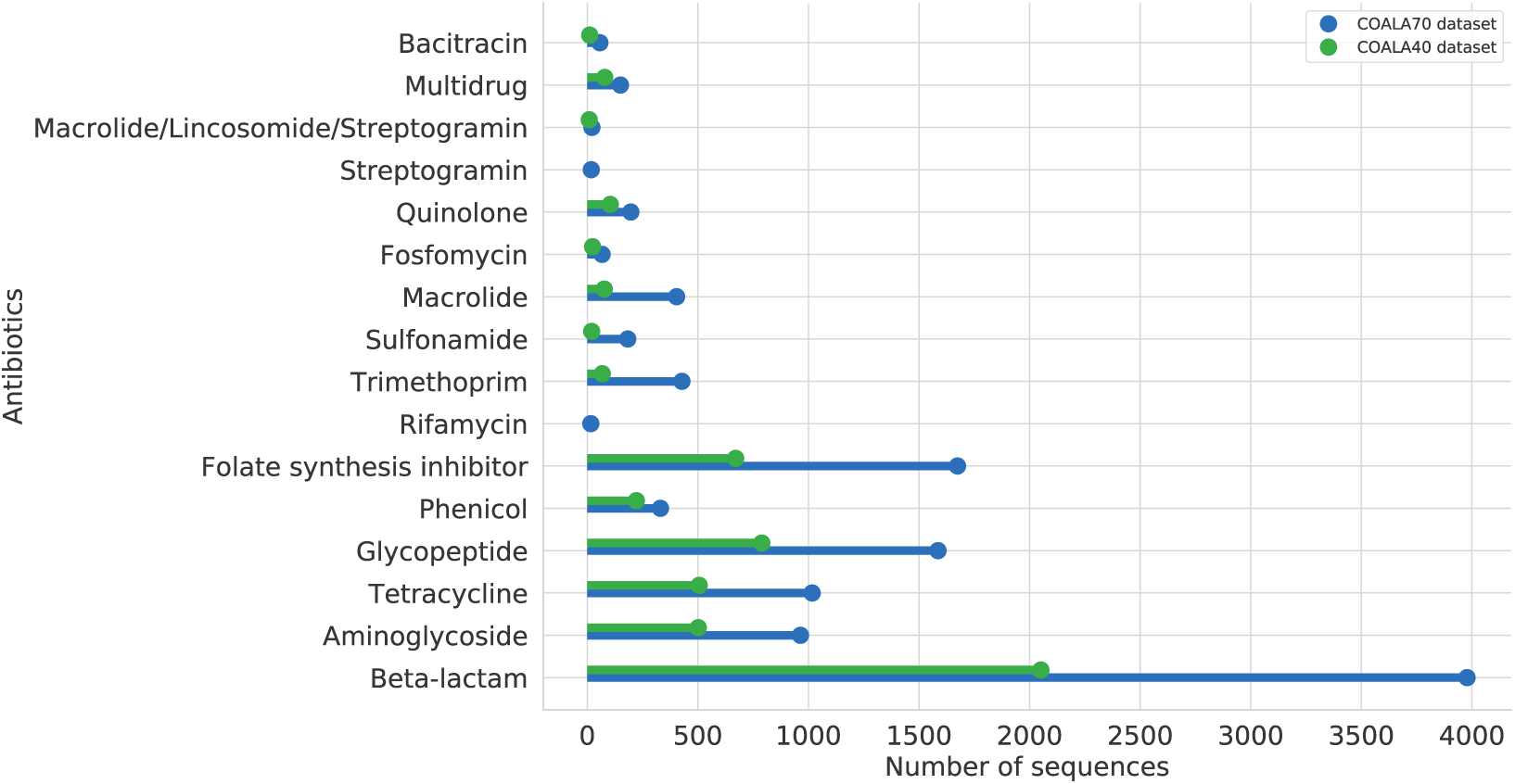
Different types of antibiotic resistance gene sequences in COALA70 and COALA40 dataset. COALA70 has 11091 sequences from 16 classes, and COALA40 has 5133 sequences from 14 classes (All numbers including for COALA100 dataset are also present in supplementary information).

## 2 Dataset

For our unlabeled dataset in the transfer learning approach, we used the UniRef50 database [25] which is comprised of ~24M protein sequences where any two sequences do not have more than 50% sequence identity. We only took the bacterial sequences (734,848) from Uniref50 as our goal is to ultimately train on a labeled dataset of antibiotic resistance bacteria protein sequences. These 734,848 sequences constitute the unlabeled dataset on which we trained a neural network as a language model.

COALA served as the labeled dataset on which we trained and tested TRAC, and other comparison methods. To construct COALA, we used 15 available antibiotic resistance gene databases mentioned in [6]. These databases are CARD [8], ResFinder [26], ResfinderFG [27], ARDB [28], MEGARes [29], NDARO [9], ARG-ANNOT [30], Mustard [31], FARME [32], SARG(v2) [10], Lahey list of β-lactamases [33], BLDB [34], LacED [35, 36], CBMAR [37], MUBII-TB-DB [38], and u-CARE [39]. We collected protein sequences from these databases with their annotations when available. All totaled, we collected 55,491 protein sequences from 58 antibiotic classes which we named COALA100, which, as its name suggests, has a high sequence redundancy rate. We used CD-HIT with two threshold levels of 70% and 40% sequence identity to produce two databases, COALA70 and COALA40 with no more than 70% and 40% sequence identity respectively. COALA70 has 11,091 protein sequences from 16 antibiotic classes, and COALA40 has 5,133 protein sequences from 14 antibiotic classes. The rationale for constructing COALA40 and COALA70 is as follows: a reliable estimate of the generalization ability of any machine learning task, it is essential that the test set does not provide instances that are identical or similar to instances in the training set. The COALA70 and COALA40 datasets therefore serve as the small labeled dataset in the second phase of transfer learning, where there are corresponding antibiotic labels for each sequence. We show all models’ performances on both datasets.

## 3 TRAC

To use transfer learning, we applied the ULMFit (Universal Language Model Fine-tuning) [18] approach used in natural language processing to perform various text classification tasks. ULMFit uses the AWD-LSTM (Averaged stochastic gradient descent Weight-Dropped LSTM) [40] architecture to train a language model on a large English language corpus. This step comprises the general-domain language model pre-training. The trained language model is then trained on the small labeled dataset on which the final text classification task will be performed, which is the language model fine-tuning. Finally, the architecture is modified to perform classification on the labeled dataset.

We used the AWD-LSTM architecture to train a neural network on the UniRef50 Bacteria sequences as a language model (Figure 2a). The architecture takes each amino acid represented as an embedding vector as input with 3 layers of LSTM [41] units. Since LSTMs (Long Short Term memory) are special recurrent neural networks that can capture long range dependency in sequential data, the AWD-LSTM model is suited to predict the next amino acid in the protein sequence. The output layer is a softmax layer where the model assigns a probability to each amino acid of being the next amino acid in the protein sequence, equivalent to the General-domain language model pretraining. Next, we trained a target task language model (Figure 2b) pretraining over the COALA40 dataset before the classification training on COALA40, the equivalent to the language model fine tuning step in ULMFit. The target task language model is trained similarly to the general domain language model in that we train the model to predict the next amino acid in a protein sequence but on the small labeled dataset which will be used for the final classification task.

**Figure 2:**
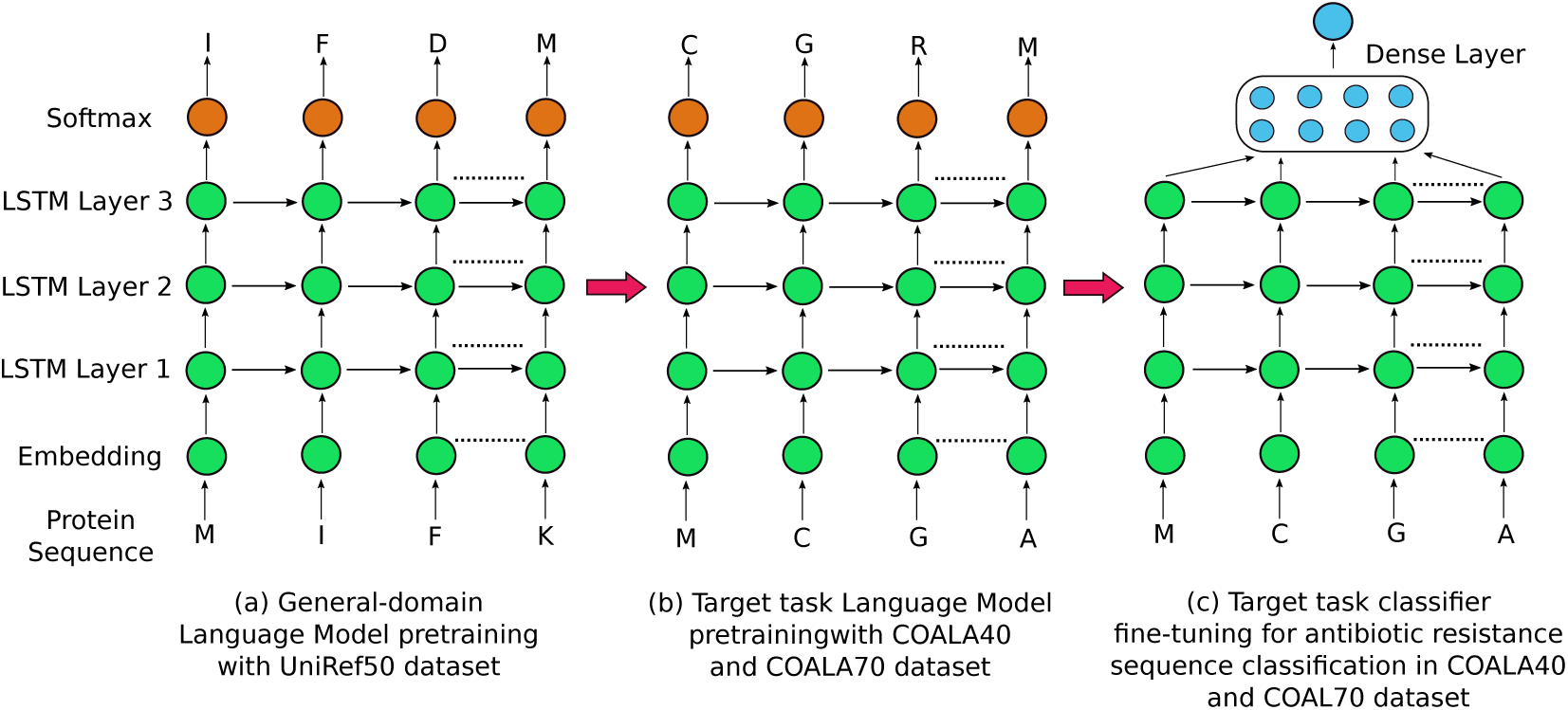
Different stages of training for TRAC. **(a)** We train the neural network on the UniRef50 dataset where the model is trying to predict the next amino acid in a protein sequence. **(b)** This is the finetuning step of the pretraining where we train the model on COALA70 and COALA40 protein sequences to predict the next amino acid. **(c)** The final classification task where the pretrained model is now trained in a supervised way to do a multiclass classification of antibiotic resistance classes.

Finally, we took this pretrained language model, and used it to classify protein sequences by antibiotics whose resistance these proteins are associated with. For both COALA40 and COALA70 datasets, we report the accuracy over a 10-fold cross validation. For both pretraining and classification, a 1-cycle learning rate schedule [42] with the AdamW [43] optimizer was used. Categorical cross-entropy [44] and label smoothing cross entropy were used for the pretaining and the classification, as respective loss functions.

## 4 Comparing Methods

We compared TRAC with other readily available AMR predicting methods that use protein sequences as input. Four current state-of-the-art methods - CARD-RGI [8], NCBI-AMRFinder [9], DeepARG [13], and SARGFAM [10] match these two criteria of availability and use of protein sequences. Besides these four methods, we developed a deep learning method that is only trained on the smaller labeled dataset as well as a random forest [45] machine learning method. This method was developed to verify whether training on an unlabeled dataset before the final classification task truly improves the performance on a labeled dataset. The random forest model was developed to verify whether deep learning and transfer learning methods perform better than traditional machine learning approaches. All six methods were tested on the COALA70 and COALA40 datasets.

### CARD-RGI

CARD-RGI integrates an antibiotic resistance ontology with homology and Single Neucleotide Polymorphism (SNP) models, and uses these models with a curated set of antimicrobial resistance sequences to identify antibiotic resistance from protein or nucleotide data.

### NCBI-AMRFinder

NCBI-AMRFinder developed from NCBI uses BLASTP [11] and HMMER [12] against the NDARO [9] database to detect antibiotic resistance from in novel protein sequences.

### SARGFAM

SARGFAM is a collection of profile HMMs built from more than 12,000 antibiotic resistance genes collected from CARD, ARDB and NCBI-NR databases. It runs an hmmsearch against these profile HMMs to detect antibiotic resistance in novel protein sequences.

### DeepARG

DeepARG is a deep learning [46] based method to classify protein sequences into the antibiotics they are resistant against. The authors created a dataset of antibiotic resistance genes from CARD, ARDB and UNIPROT. Sequences from UNIPROT were aligned against CARD and ARDB with Diamond [47], and the normalized bit scores from the alignment were used as features for the deep learning model.

### Baseline Deep Learning

To develop a deep learning model trained on the COALA70 and COALA40 datasets, we used a self-attention based sentence embedding model introduced in [48]. For input, we represented each amino acid in a protein sequence as an embedding vector which was randomly initialized, and then trained end-to-end. We used a Long Short Term Memory or LSTM [41] recurrent neural network which takes these embedding vectors as input. Following that is the self-attention part, essentially a feed-forward neural network with one hidden layer. This network takes the output from the LSTM layer as input, and outputs weights which are known as attentions. We multiplied the outputs of the LSTM layer by these attentional weights to get a weighted view of the LSTM hidden states. The product then is passed to the next layer of the network, and the final layer is a softmax layer which assigns a probability to each class of antibiotics so that they sum to 1. In training, AdamW [43] optimizer with a 1-cycle learning rate schedule [42] with a negative loss likelihood loss function was used.

### Baseline Random Forest

We also applied a random forest model on the COALA70 and COALA40 datasets, and assessed its performance. For features, we used 3-mer count for each protein sequence. The hyper-parameters of the random forest model were tuned with random search using the Scikitlearn [49] machine learning library.

## 5 Results

Figure 4 shows the comparison between all the models with the COALA40 and COALA70 datasets. We use accuracy to compare the models. For a set of predictions *Ŷ* = (*ŷ*_1_, *ŷ*_3_,…*ŷ*_*N*_) and a set of true labels *Y* = (*y*_1_, *y*_3_,…*y*_*N*_) accuracy is defined as:

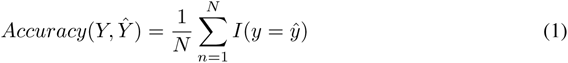

where *I* is the indicator function defined as:

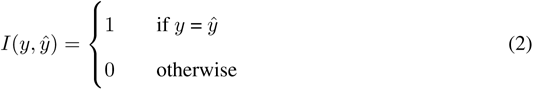

**Figure 3:**
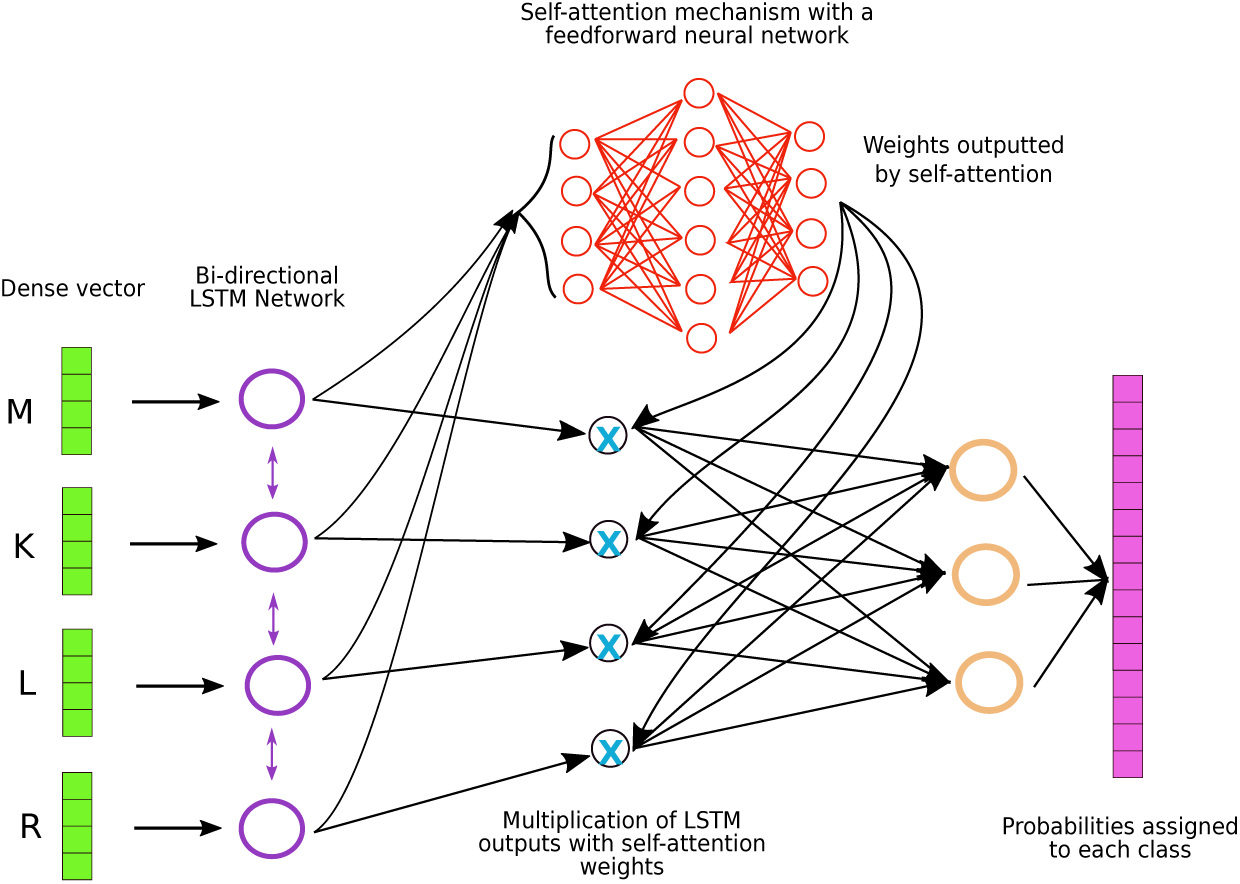
The baseline model for antibiotic resistant gene classification. Each amino acid in a protein sequence is represented by a vector. These vectors are used as input for LSTM neural networks [41]. The output of the LSTM network goes into a simple feed-forward network, and the output of this feed-forward network is weights which sum to 1. These weights are multiplied with the outputs of the LSTM network to give them weights. This mechanism lets the model learn which input from the protein sequence is crucial in classifying a gene into its correct antibiotic class. Finally, there is a softmax layer, the size of which is 15 corresponding to the 15 antibiotic classes. This layer assigns a probability to each class, the class that is assigned the highest probability becomes the prediction for a certain protein sequence.

**Figure 4:**
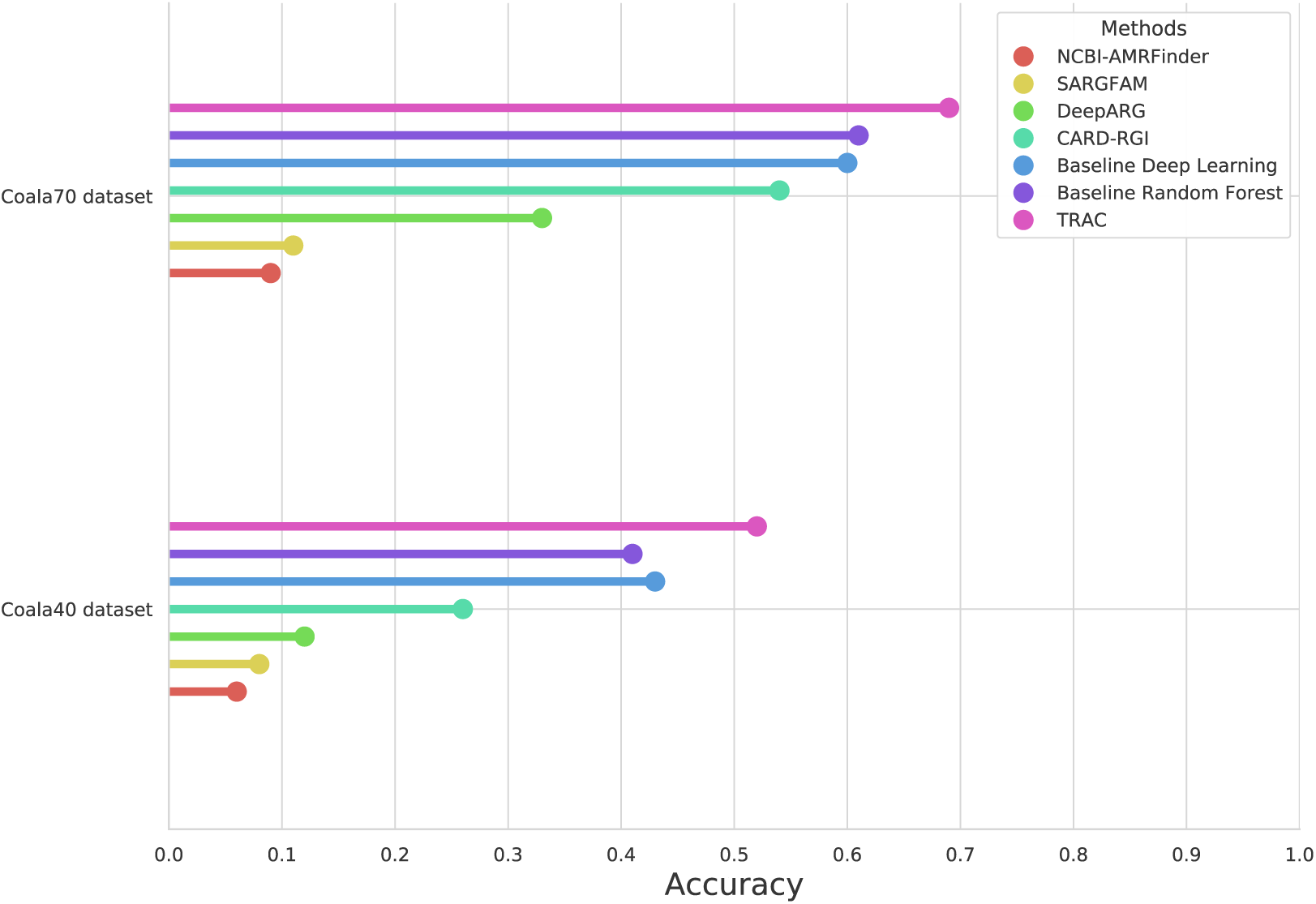
Accuracies of all methods tested on both COALA70 and COALA40 datasets.

The accuracy is the mean of 10 × cross validation on both COALA40 and COALA70 datasets. The hyperparameters were chosen by nested cross validation so that we do not overestimate on the final test set. For the COALA40 dataset, we see that TRAC outperforms all other models with an accuracy of 0.52, and the next closest performing model is our own deep learning model that we constructed without any pretraining (accuracy 0.43). Random forest performs almost with a similar accuracy of 0.41. The other methods perform not as well: CARD-RGI (accuracy 0.26), DeepARG (accuracy 0.12), SARGFAM (accuracy 0.08) and NCBI AMRFinder (accuracy 0.06). In the COALA70 dataset, TRAC again outperforms all other methods by a large margin (accuracy 0.69). Interestingly, a well tuned random forest model (accuracy 0.61) performs similarly to a deep learning model (accuracy 0.60) that was not pretrained. With a bigger dataset of COALA70, performance of CARD-RGI increases (accuracy 0.54). In contrast, DeepARG, SARGFAM and NCBI AMRFinder still lag behind. The performance of all the models on both COALA40 and COALA70 dataset can also be seen in Table 1 in Supplementary Materials.

## 6 Discussion

Identifying genes that confer resistance to antibiotics is crucial to understand what types of resistance exist in a microbial sample. In this study, we compare currently available computational tools to predict the antibiotic class that to which a gene might confer resistance. Towards this end, we developed a dataset, COALA, that contains protein sequences from 15 antibiotic resistance databases available online. We also curated the antibiotic resistance class labels for these protein sequences from their respective databases. This resulted in a dataset that has huge number of redundant and similar sequences. To gain reliable estimate of how well all the computational methods perform, we removed sequence redundancies to 40% and 70% identity thresholds, resulting in two datasets, COALA40 and COALA70. By doing so, especially with COALA40, we remove most of the homology-detection component from the

We can see from the performances of all models that TRAC, the transfer learning approach performs is considerably more accurate. Even so, TRAC gains around 69% accuracy for COALA70 and around 52% accuracy for COALA40. This indicates that multiclass classification for antibiotic resistance class prediction is indeed a difficult task. However, the alignment free approaches which are TRAC, Deep learning without pretraining, and random forest all perform better than the other four methods: NCBI-AMRFinder, SARGFAM, CARD-RGI, and DeepARG which depend, to different extents, on alignment-based approaches such as BLAST or HMMER. This is crucial with respect to the fact that performing CD-HIT with such low thresholds of identity framed this problem as a remote homolog prediction problem. Within the alignment free approaches, surprisingly, a deep learning model without pretraining did not perform better than a well-tuned random forest model that was trained on *k*-mer features. This leads to the question of why TRAC is performing significantly better than deep learning without pretraining. We hypothesize that by training on unlabeled bacterial sequences and trying to predict the next amino acid in a sequence, TRAC gains an internal representation of bacterial sequences which helps it to identify remote homologs in the downstream classification task. Another possible reason is the labeled datasets, especially COALA40, do not have enough training sequences for a deep learning model to learn without pretraining. TRAC mitigates this problem by shifting the burden of learning onto the unlabeled dataset for pretraining.

We hope that the COALA70 and COALA40 datasets will provide good test beds for developing novel machine learning methods for AMR prediction with superior accuracy performance equivalent to that provided by the MNIST dataset [50] for the deep learning community. The performance of TRAC provides further proof that language model pretraining boosts model performance with few labeled data [51] which is the case in many bioinformatics tasks. For future work, it will be interesting to unpack the internal representation of the protein sequences learnt by TRAC, and how it compares with traditional bioinformatics features such as Position Specific Scoring Matrices (PSSM) or HMMs. We are also excited with the prospects of integrating the latest advancements in the Natural language field such as Transformer [52] inspired encoding techniques (e.g. BERT [53]) into the pretraining step, and improving antibiotic resistance class prediction adding to current clinical practices.

