## Supplementary Material for "Transfer learning improves antibiotic resistance class prediction"

### Terminal commands

CD-HIT command to make the COALA70 dataset from COALA100 -

```
$ cd-hit -i coala_db.fa -o coala_db_cd_hit_0.7.fa -c 0.7
```

CD-HIT command to make the COALA40 dataset from COALA100 -

```
$ cd-hit -i coala_db.fa -o coala_db_cd_hit_0.4.fa -c 0.4 -n 2
```

**Table 1:** Mean accuracy from 10-fold nested cross validation for all methods over both COALA40 and COALA70 datasets. Standard deviation of accuracy is shown for our models where cross validation is done.

| | **COALA40 dataset** | **COALA70 dataset** |
|---|---|---|
| CARD-RGI | 0.261 | 0.544 |
| NCBI-AMRFinder | 0.062 | 0.093 |
| SARGFAM | 0.085 | 0.110 |
| DeepARG | 0.126 | 0.330 |
| Baseline Random Forest | 0.416 ± 0.007 | 0.614 ± 0.065 |
| Baseline Deep Learning | 0.436 ± 0.018 | 0.609 ± 0.097 |
| TRAC | **0.520 ± 0.084** | **0.696 ± 0.078** |

**Table 2:** Antibiotic and the number of respective resistance sequences present in the COALA100 dataset.

| Antibiotic | Number of sequences |
|---|---|
| BETA-LACTAM | 29099 |
| MULTIDRUG | 4814 |
| TETRACYCLINE | 4224 |
| FOLATE-SYNTHESIS-INHABITOR | 3429 |
| GLYCOPEPTIDE | 3124 |
| AMINOGLYCOSIDE | 2245 |
| TRIMETHOPRIM | 1764 |
| MACROLIDE/LINCOSAMIDE/STREPTOGRAMIN | 1693 |
| PHENICOL | 1469 |
| MACROLIDE | 1015 |
| FOSFOMYCIN | 573 |
| SULFONAMIDE | 560 |
| QUINOLONE | 498 |
| BACITRACIN | 295 |
| POLYMYXIN | 249 |
| RIFAMYCIN | 64 |
| COLISTIN | 54 |
| PHENICOL/QUINOLONE | 43 |
| LINCOSAMIDE | 38 |
| STREPTOGRAMIN | 36 |
| QUATERNARY AMMONIUM | 33 |
| FOSMIDOMYCIN | 29 |
| BLEOMYCIN | 18 |
| LINCOSAMIDE/STREPTOGRAMIN | 15 |
| MACROLIDE/STREPTOGRAMIN | 14 |
| STREPTOTHRICIN | 12 |
| FUSIDIC ACID | 11 |
| TETRACENOMYCIN | 9 |
| NITROIMIDAZOLE | 7 |
| AMINOGLYCOSIDE/QUINOLONE | 6 |
| PENICILLIN | 4 |
| ACRIFLAVINE/AMINOGLYCOSIDE/BETA-LACTAM/GLYCYLCYCLINE/MACROLIDE | 4 |
| FLUOROQUINOLONE | 4 |
| MUPIROCIN | 3 |
| FUSARIC-ACID | 3 |
| AMPICILLIN/STREPTOMYCIN/TETRACYCLINE/SULFONAMIDE | 3 |
| PUROMYCIN | 3 |
| ACRIFLAVINE | 2 |
| THIOSTREPTON | 2 |
| PLEUROMUTILIN | 2 |
| TELLURITE | 2 |
| AMINOCOUMARIN | 2 |
| TUBERACTINOMYCIN | 2 |
| AMINOPTERIN/TRIMETHOPRIM | 1 |
| AVILAMYCIN | 1 |
| MACROLIDE/PLEUROMUTILIN | 1 |
| AMPICILLIN/BETA-LACTAM/PENICILLIN | 1 |
| BICYCLOMYCIN | 1 |
| CEPHALOTHIN/AMPICILLIN | 1 |
| CANAVANINE | 1 |
| COLICIN I | 1 |
| COLICIN | 1 |
| GALACTOSE | 1 |
| MOLYBDENATE | 1 |
| ISONIAZID | 1 |
| PENICILLIN G/NORFLOXACIN/CHLORAMPHENICOL/GENTAMICIN | 1 |
| SERINE | 1 |
| PHENICOL/OXAZOLIDINONE | 1 |

**Table 3:** Antibiotic and the number of respective resistance sequences present in the COALA70 and COALA40 dataset. Some categories are absent in the COALA40 dataset.

| | **Number of sequences** | |
|---|---|---|
| **Antibiotic** | COALA70 | COALA40 |
| BETA-LACTAM | 3978 | 2051 |
| FOLATE-SYNTHESIS-INHABITOR | 1674 | 671 |
| GLYCOPEPTIDE | 1586 | 789 |
| TETRACYCLINE | 1017 | 506 |
| AMINOGLYCOSIDE | 964 | 502 |
| TRIMETHOPRIM | 428 | 68 |
| MACROLIDE | 404 | 77 |
| PHENICOL | 331 | 0 |
| QUINOLONE | 197 | 104 |
| SULFONAMIDE | 183 | 20 |
| MULTIDRUG | 150 | 79 |
| FOSFOMYCIN | 67 | 24 |
| BACITRACIN | 57 | 11 |
| MACROLIDE/LINCOSAMIDE/STREPTOGRAMIN | 21 | 9 |
| STREPTOGRAMIN | 18 | 0 |
| RIFAMYCIN | 16 | 0 |
